# *De novo* assembly and annotation of the singing mouse genome

**DOI:** 10.1101/2022.07.29.502048

**Authors:** Samantha K. Smith, Paul W. Frazel, Alireza Khodadadi-Jamayran, Paul Zappile, Christian Marier, Mariam Okhovat, Stuart Brown, Michael A. Long, Adriana Heguy, Steven M Phelps

## Abstract

**Background:** Developing genomic resources for a diverse range of species is an important step towards understanding the mechanisms underlying complex traits.Specifically, organisms that exhibit unique, accessible phenotypes-of-interests, allow researchers to address questions that may be ill-suited to traditional model organisms. We sequenced the genome and transcriptome of Alston’s singing mouse (*Scotinomys teguina*), an emerging model for social cognition and vocal communication. In addition to producing advertisement songs used for mate attraction and male-male competition, these rodents are diurnal, live at high-altitudes, and are obligate insectivores, providing opportunities to explore diverse physiological, ecological, and evolutionary questions.

**Results:** Using PromethION, Illumina, and PacBio sequencing, we produced an annotated genome and transcriptome, which were validated using gene expression and functional enrichment analyses. To assess the usefulness of our assemblies, we performed single nuclei sequencing on cells of the orofacial motor cortex, a brain region implicated in song coordination, identifying 12 cell types.

**Conclusions:** These resources will provide the opportunity to identify the molecular basis of complex traits in singing mice as well as to contribute data that can be used for large-scale comparative analyses.

## Background

The rapid development of sequencing tools in the last 20 years has allowed interrogation of coding and noncoding sequence evolution (1–6), gene regulation (7– 15), protein-genome interactions (16–19), and many other processes (20–22). Although initial focus was on a few model organisms, genomic sequencing and analysis has become increasingly feasible for nontraditional species. This ability to characterize the genomes of nontraditional organisms has created opportunities to assess historically challenging questions from across biological disciplines.

Nontraditional rodents are particularly useful models for understanding the mechanisms underlying complex traits. For example, different species of deer mice construct burrows of varying complexity, a trait that has a strong genetic component (23). Nontraditional rodents are particularly tractable since many of the tools developed in laboratory mice and rats are easily adapted to related species. The translational nature of these powerful tools provides avenues for examining traits that are lacking in inbred, laboratory rodents. Finally, the extensive mapping of the rodent brain and the availability of tools for measuring and manipulating neural activity provide a strong basis for understanding the neurobiology of complex traits. Although such tools enable mechanistic work, it is also important to develop genomic resources for nontraditional rodents, as the examination of certain mechanisms (e.g., DNA methylation) are best facilitated by the availability of an annotated genome (24).

Alston’s singing mouse, *Scotinomys teguina*, produces a unique, easily quantifiable vocal display that makes it an excellent model for understanding the genomic mechanisms of complex, behavioral traits. These diurnal rodents live in the montane grasslands of central America and are obligate insectivores (25). Singing mice are named for the long, elaborate songs they use for mate attraction and male-male competition (25–29). Their unique natural history as well as their complex social interactions make singing mice an excellent candidate for exploring the mechanisms and evolution of traits such as circadian rhythms, diet and energy balance, the challenges of thermoregulation or high-altitude living, dynamic vocal communication, and more.

Unlike model rodents such as house mice, singing mice produce highly structured advertisement songs (Figure 1) which make them an emerging model for social cognition and vocal communication. These songs consist of rapidly repeated frequency-modulated notes which span ∼16 kHz in as little as 12 msec (30). Note amplitudes, frequencies, and repetition rates are modulated over the course of the song (Figure 1). Singing mice have highly structured vocalizations that are rapidly exchanged with conspecifics (counter-singing) in a manner whose timescale resemble human conversational speech, a feature not found in house mouse communication (31).

**Figure 1.**
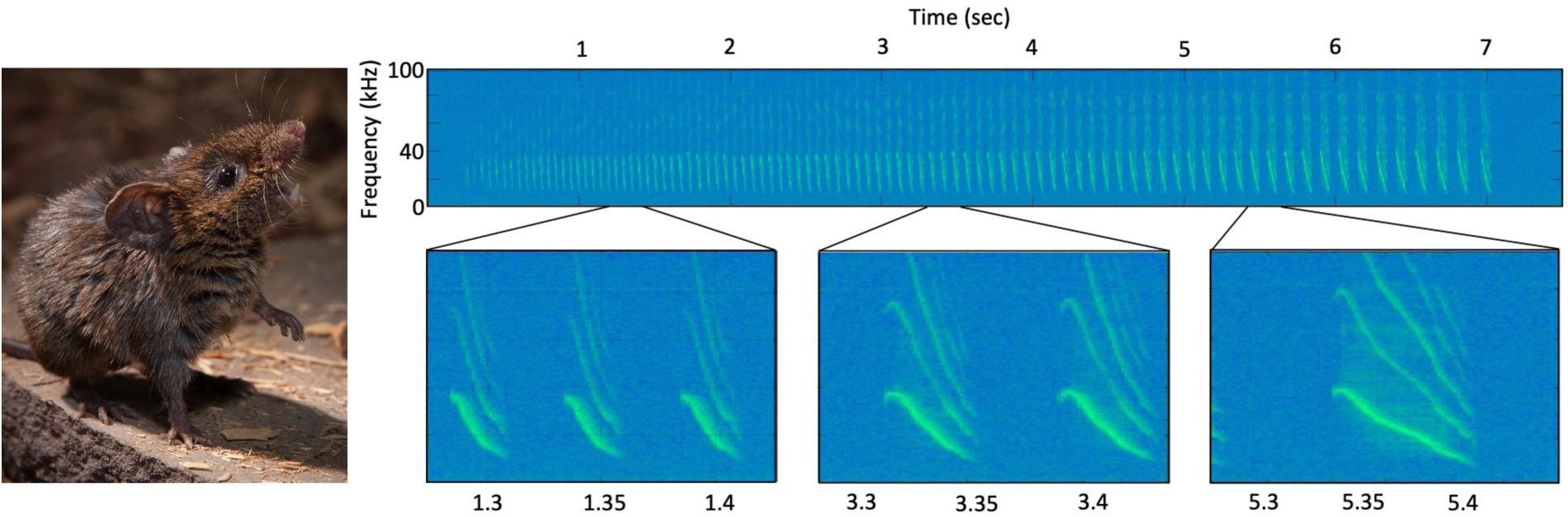
(Left) A singing mouse and (right) a spectrogram of a representative advertisement song. Insets below show how frequency bandwidth, note shape, and note length change over the course of a song. Photo: Long lab. Figure adapted from (34).

In addition to variation among species (27,30), the advertisement song also varies between individuals (32) and populations of singing mice (30). Among individuals, both internal and external cues drive song differences. For example, androgen levels and adiposity signals such as circulating leptin are associated with differences in “song effort” measures (e.g., song length), but not spectral features (e.g., frequency bandwidth) which may be set by vocal anatomy (28,29,32–34). The social environment also tunes vocal output. For example, when other males are present, male singing mice produce longer songs and rapidly turn-take during singing bouts, finely coordinating song onset and offset (31). Together, these unique song features and the complexity of cues that impact song provide an exemplary opportunity to understand many fundamental questions such as how internal and external cues are integrated to modulate behavior, how elaborate vocalizations evolve, and more. The development of genomic resources for singing mice will provide opportunities to explore these questions as well as contribute to broad, comparative work.

We sequenced the singing mouse genome and transcriptome using PromethION, Illumina, and PacBio technologies. We next assembled and annotated the genome and transcriptome and examined gene expression to assess the utility of our assemblies. Finally, we extracted and sequenced cell nuclei from the orofacial motor cortex (OMC) using 10X Genomics. The OMC was chosen because it plays an important role in the social modulation of singing behavior, namely counter singing between conspecifics (31). To identify cell-types within the OMC, we used gene expression analysis. The annotated genome and transcriptome data are available on the UCSC genome browser (https://bit.ly/3hfUiIy). Single-nuclei OMC data are available upon request. A well-annotated genome and transcriptome will enable future work identifying the genomic substrates of a variety of physiological and behavioral adaptations.

## Methods

### DNA isolation

All animal procedures were approved by the University of Texas at Austin and New York University Grossman School of Medicine IACUC. For PacBio and Illumina sequencing, we sacrificed 1 adult, male singing mouse. Liver and brain were dissected out and flash-frozen immediately. GDNA was then extracted from tissues using a Qiagen DNeasy kit. We visualized DNA integrity on an agarose gel and quantified DNA quality using a nanodrop. For PromethION sequencing, we sacrificed an adult, male singing mouse, extracted its liver, and immediately froze the tissue. High molecular weight DNA was extracted from liver tissue using a Genomic-tip 20/G DNA kit (QIAGEN, 10223).

### DNA sequencing and genome assembly

We did library preparations and sequencing using PromethION technology at the New York University Langone Medical Center.

PacBio library preparation and sequencing was done at Duke University using 6-8 kb insert sizes with sub-reads ranging from 2KB-3KB. We did Illumina library preparation and sequencing at the University of Texas Austin Core facility. Two Illumina libraries were created: a fragmentation library consisting of 170, 400, and 900 bp segments (PE Barcode+2×100, 3 lanes = 510M reads requested) and a mate-pair library with a 3KB insert size (PE Barcode+2×100, 1 lane=170M reads requested).

We assembled long reads from PacBio and PromethION and short reads from 10X genomics using the mixed read assembly tool MaSuRCA (v. 3.2.8) (35). The assembled reference genome was masked for repeats using RepeatMasker (v. 1.332) (36).

### RNA isolation and sequencing

RNA extraction and sequencing was done at UT Austin and NYULMC. All animals were sacrificed using isoflurane overdose. At UT Austin, forebrain, hindlimb skeletal muscle, gonads, and liver were dissected from 1 adult male singing mouse and immediately flash frozen. We extracted total RNA using a standard TRIzol method. RNA was then submitted to the UT Core facility for library preparation and Illumina sequencing. For RNAseq performed at NYU, we extracted RNA from freshly dissected tissue of two male and two female singing mice (liver and brain) using a Qiagen RNeasy Mini kit (Qiagen 74104). We then homogenized the tissue using a rotor-stator homogenizer with disposable tips and did on-column DNAse digestion following manufacturer’s instructions. An automated system performed poly-A library prep, and samples were run on a single-read Illumina HiSeq 4000 flowcell.

### Transcriptome assembly, and analysis

#### Transcriptome assembly

We assembled a *de novo* and reference guided transcriptome using Trinity (v2.8.4) (37,38). For the guided transcript assembly all the RNA-Seq reads were mapped to the assembled reference genome using STAR mapper (v2.5.0c) (39). Alignments were guided by a Gene Transfer Format (GTF) file. For quality control, we mapped the RNA reads to the assembled transcripts provided by Trinity (37,38). More than 83% of the reads mapped properly, suggesting a high-quality transcript assembly.

#### Differential expression analysis

We calculated the mean read insert sizes and their standard deviations using Picard tools (v. 1.126) (40). Read count tables were generated using HTSeq (v0.6.0) (41) and normalized based on library size factors using DEseq2 (42). For differential expression analysis, we used BEDTools (v2.17.0) (43) and bedGraphToBigWig tool (v. 4; ENCODE) (44,45) to generate read-per-million (RPM) normalized BigWig files. To compare gene expression across samples and their replicates, we used principal component analysis and Euclidean distance-based sample clustering. All downstream statistical analyses and plot generation were performed in R (v3.1.1) (46).

#### Functional enrichment

*GO and KEGG analyses*. To assess the accuracy of transcriptome assembly and annotation, GO MWU (47) and KEGG (48–50) analyses were used. The GO MWU method of gene ontology (GO) enrichment analysis uses a ranked list of genes to identify whether each GO category is significantly enriched with up-or down-regulated genes (47,51). We did functional enrichment analysis of GO and KEGG Reactome pathways using g:Profiler (v. e101_eg48_p14_baf17f0) with a g:SCS significance threshold of 0.05 (52). Ordered gene lists for each tissue type included only those that had a |fold-change| of at least 2. We exported GO functional enrichment results from g:Profiler and created network pathways (53) using the EnrichmentMap application (54) in Cytoscape (55). Maps were created with FDR *Q* value <0.01 and combined coefficients >0.375 with a combined constant of 0.5. We used an expression file of normalized fold-change values to create heatmaps of genes enriched pathways. We then used AutoAnnotate to interpret the function of groups of nodes in the network.

### Genome annotation

RNA-Seq reads from Illumina and the assembled reference genome were used to create transcript-backed and prediction-based annotations. We concatenated both the guided and *de novo* transcriptome assembly results and cleaned them using the PASA pipeline (v.) (56) for UniVec vector sequences (57). Cufflinks (58–61) was used to make a GTF file for PASA pipeline and the tdn.accs file was made using the *de novo* assembly. We used the GTF file produced by PASA’s comprehensive pipeline as our reference GTF for downstream analysis (bulk RNA-Seq and scRNA-Seq). A cDNA fasta file was produced from the GTF and used as an input for BLAST (62). We blasted the cDNA file against the Uniprot database (63). BLAST results were then used to annotate the GTF file with gene symbols.

### Single nuclei sequencing and analysis

#### Single nuclei sequencing

We tagged the orofacial motor cortex (OMC) area from one adult male singing mouse for extraction via stereotaxic injection of fluorescent dextran beads into the brain as previously described (31). Post injection, the mouse was immediately transcardially perfused with ice-cold artificial cerebrospinal fluid (aCSF).

We sectioned the brain into 250 µm sections, located the dyed area under a dissecting microscope, and removed the region with a scalpel. Extracted tissue was immediately flash frozen in liquid nitrogen and stored overnight at -80°C. We dissociated nuclei using a modified version of the Mccarroll lab protocol (64). FITC-tagged NeuN antibody (Sigma, MAB377) was prepared following manufacturer’s instructions (Abcam, 188285), and used to enrich for NeuN+, DAPI+ nuclei on a MoFlo XDP flow cytometer (Beckman Coulter) with a 100 µm nozzle. We loaded 9000 sorted nuclei into GEMs on a 10X Genomics Chromium Controller (1^st^ generation, 10X v3 chemistry) using 3’ v3 chemistry and recovered 3500 high-quality nuclei after standard analysis (10X Genomics CellRanger pipeline v. 3.1.0) (65).

#### Single nuclei gene expression analysis

Single-nuclei gene expression data were generated using the 10X Genetics Chromium system, following the manufacturer’s instructions for sample and library prep. We aligned raw FASTQ files to the singing mouse transcriptome and then assigned reads to individual nuclei via the 10X CellRanger pipeline. The resulting gene expression matrix was analyzed using the standard Seurat package (v. 3) (66) in RStudio (v. 4.0.2). We excluded genes with expression in <3 nuclei from the analysis. We filtered the expression data to only keep nuclei with fewer than 11,500 genes, and fewer than 40,000 molecules detected, excluding 14 likely doublet nuclei.

Singlet nuclei gene expression data were then log-normalized using the Seurat pipeline (66) and only the top 2,000 most variable genes were selected for downstream analysis. We ran PCA on the top 2,000 variable genes using standard Seurat settings and clustered nuclei via standard commands using the first 20 principal components. A resolution value of 0.1 was used to capture large, cell-type-level clusters of similar nuclei, resulting in 12 clusters that were categorized into major cell types based on known marker genes. We did dimensional reduction via two standard methods, tSNE (t-distributed Stochastic Neighbor Embedding) (67) and UMAP (Uniform Manifold Approximation and Projection) (68,69). UMAP better distinguished clusters and was used for downstream analyses. Marker genes for each cluster (genes significantly enriched) were identified using the standard threshold values of >0.25 percent of nuclei expressing the gene and >0.25 log-fold change. We plotted the top 10 markers genes for each cluster on a heatmap and compared these with known marker genes to determine what cell types are represented by each cluster.

## Results

The annotated singing mouse genome and transcriptome can be accessed at the UCSC genome browser (https://bit.ly/3hfUiIy).

### Genome and transcriptome assembly and annotation

After assembly, the total genome size was 2.4 billion base pairs. After scaffolding there were 7806 contigs. We assembled both a *de novo* and reference guided transcriptome and identified 754,907 transcripts. Contig N50 from the *de novo* assembly was 826 for all transcript contigs (median contig length: 351, average contig: 620.03) and 513 when considering only the longest isoform per gene (median contig length: 327, average contig: 478.26). We found an 83.41% overall alignment rate of the transcriptome to the genome.

We annotated the genome using the PASA pipeline and resulting GTF file can be accessed at the UCSC genome browser (https://bit.ly/3hfUiIy). We annotated genes using blast and only included annotations for genes that had at least 80% sequence similarity to the reference gene (14,989 genes included).

### Validation

We validated the quality of the transcriptome and annotations by doing gene expression analyses. Reads were normalized using DEseq2 (42) and samples were clustered by Euclidean distance (Figure 2). As expected, samples clustered by tissue type. Principal components analysis revealed two components that distinguished tissue type (Figure 3). PC1 separated brain gene expression from that of the liver, while PC2 distinguished brain and liver expression from that of the muscle and gonads. We then compared gene expression profiles between pairs of tissues.

**Figure 2.**
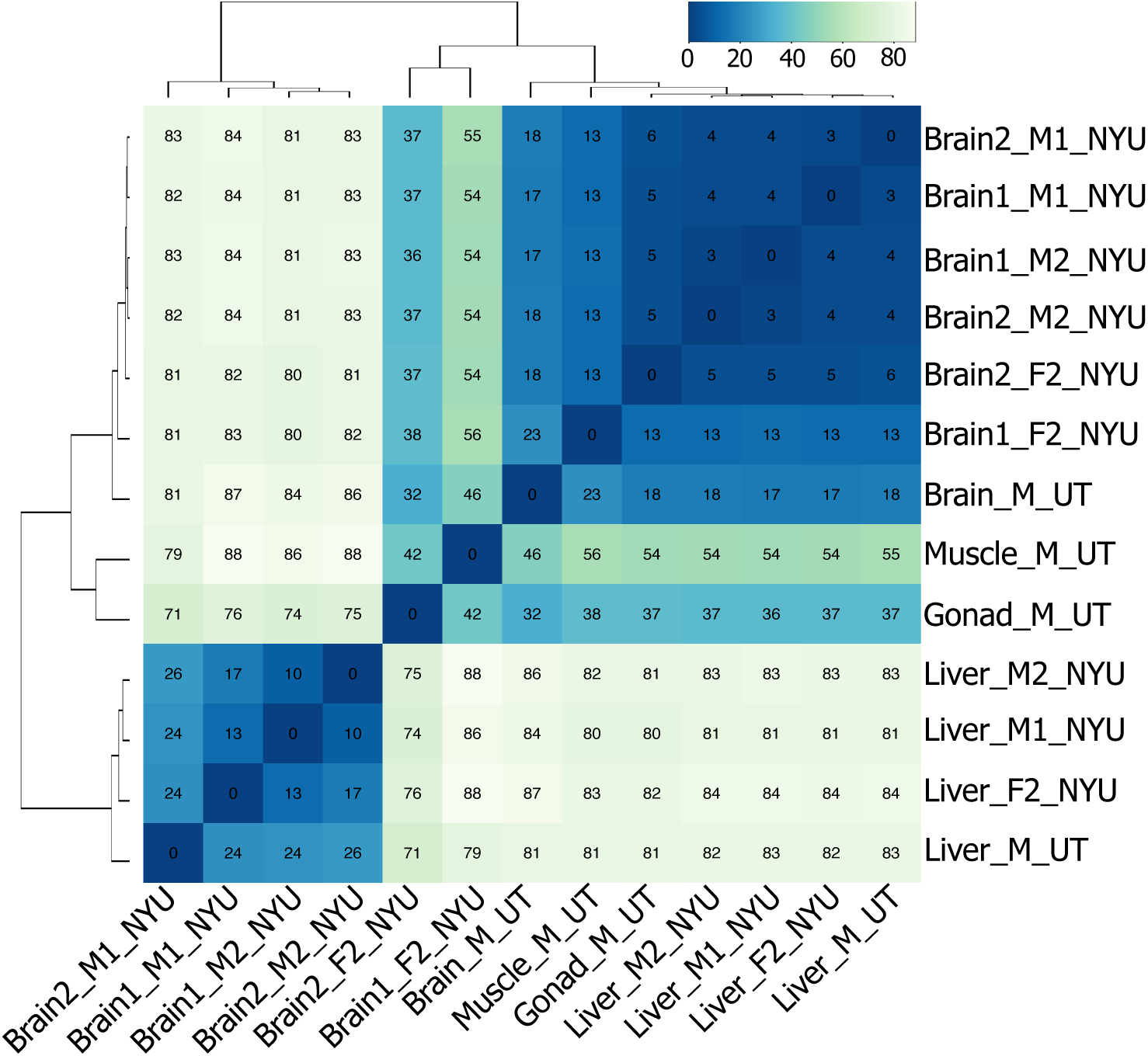
Euclidean-distance-based heatmap shows that samples of the same tissue type have the most similar gene expression. Lower values (darker blue) indicate more similarity.

**Figure 3.**
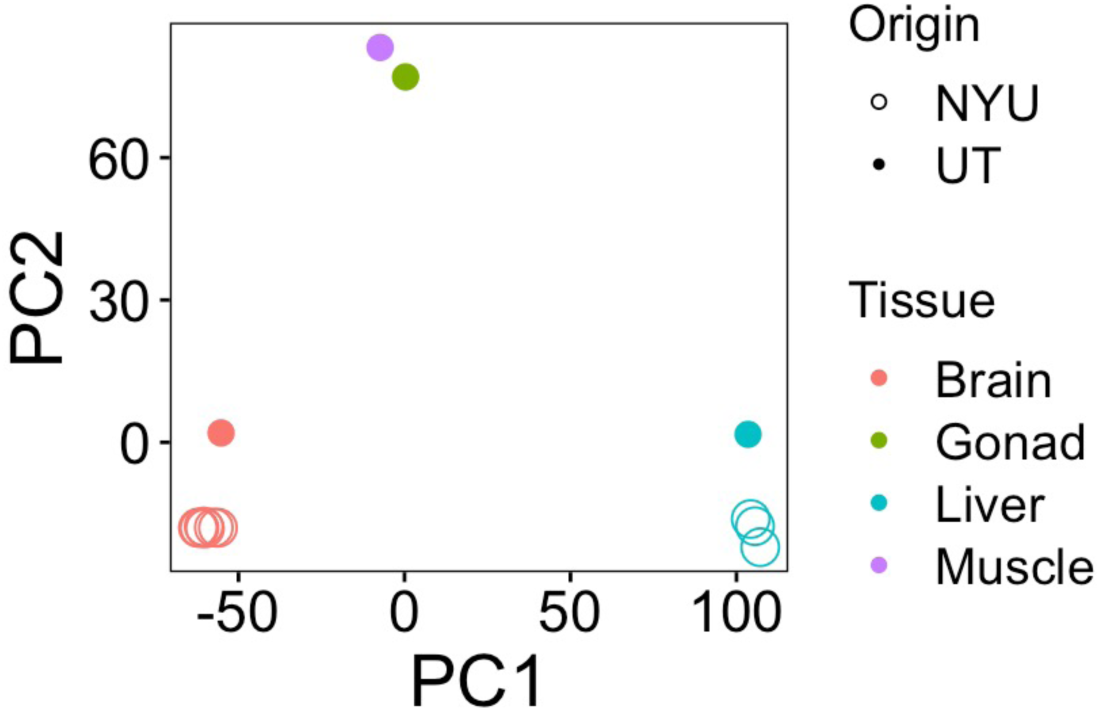
Biplot of the first two principal components, which distinguish tissue types. Brain and liver gene expression drive PC1, while the differences between brain/liver gene expression and that of the gonad and liver underlie PC2. Dot color indicates tissue type and whether the circle is filled in or not indicates where the samples were collected.

To validate that we accurately mapped transcripts to annotated genes, we did GO MWU (47) and KEGG (48–50) analyses. We found that the metabolic pathways KEGG term was the most significantly enriched among all annotated genes within the genome (Figure 4). We then did GO MWU (47) on each tissue-type gene list which we ranked by fold-change. This analysis revealed enrichment of expected GO terms based on tissue type. For example, genes upregulated in the brain were enriched with terms related to synapse structure and function (Figure 5a). To further assess whether we detect of appropriate, tissue-specific gene expression, we constructed network pathways from the brain GO enrichment results. The analysis determined 389 gene sets (“nodes”) and 770 instances of overlap between gene sets (“edges”) that were sorted into 146 clusters (Figure 5b). We found that the network was annotated with functions consistent with first-principles predictions based on the focal tissue. For example, the brain functional GO network was annotated with functions such as “postsynaptic membrane component”.

**Figure 4.**
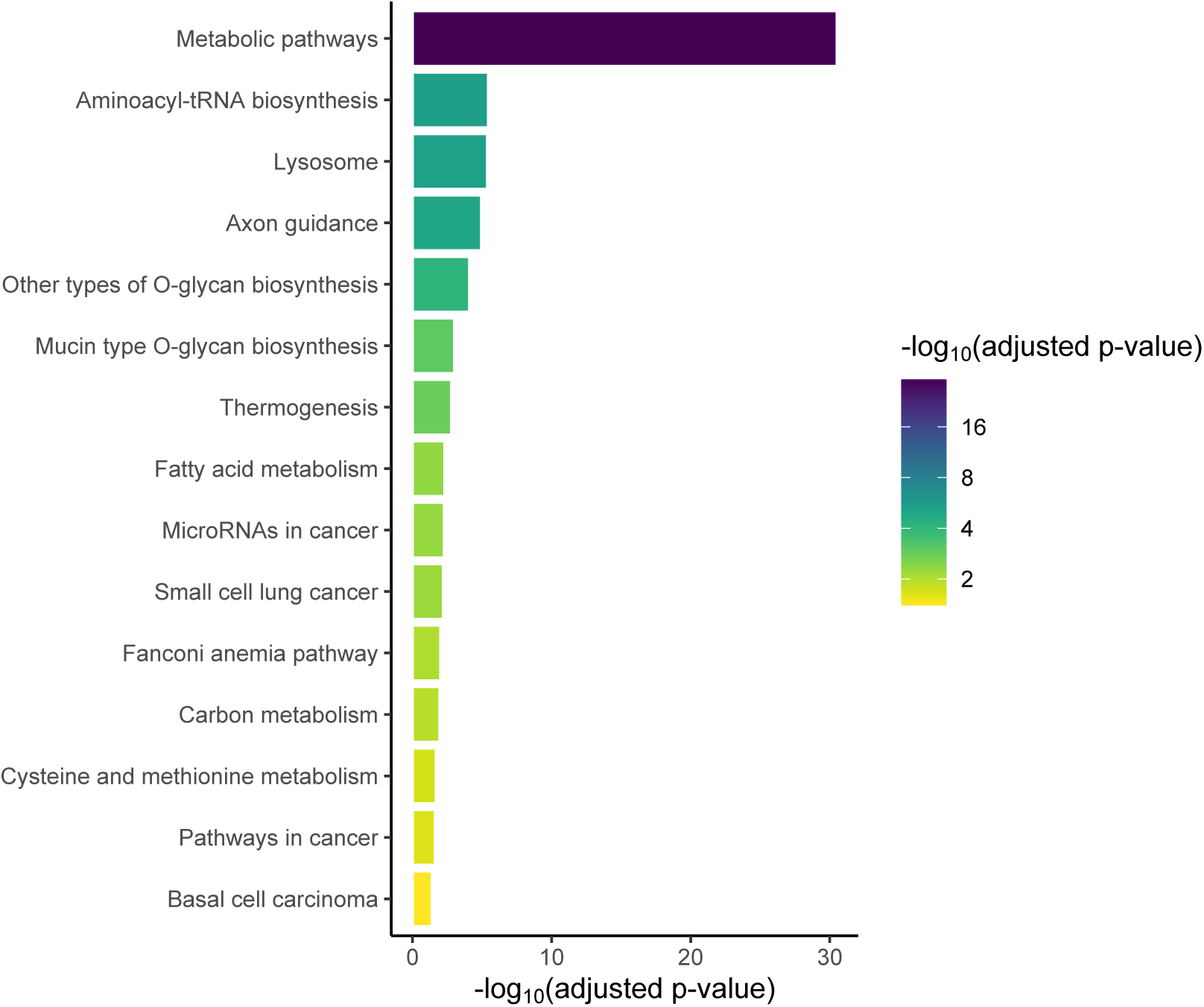
Barplot of KEGG terms for all annotated genes shows that metabolic pathways are enriched in this dataset.

**Figure 5.**
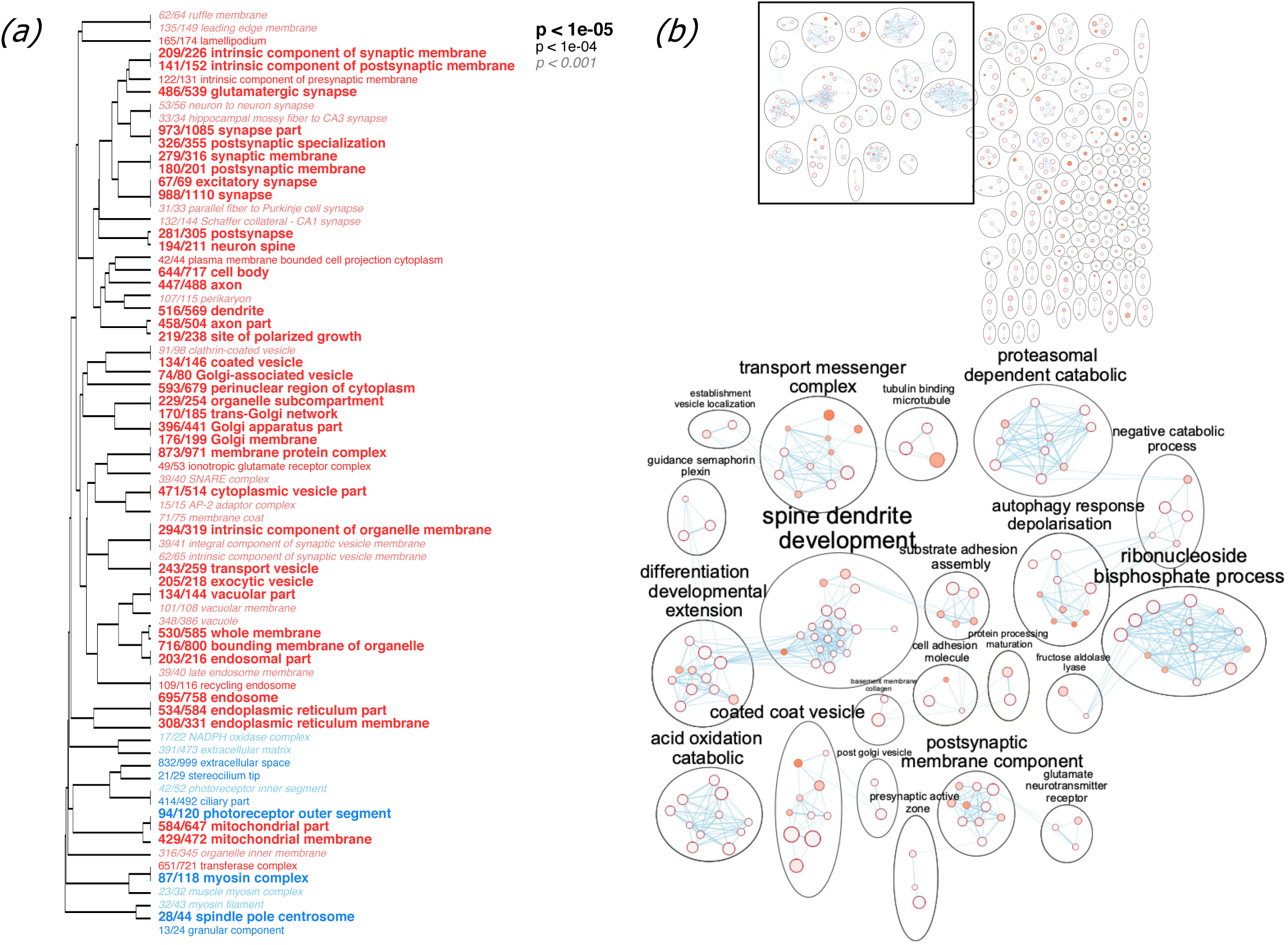
(a) GO tree of enriched terms for ranked brain gene expression (GO division: cellular compartment) using Fisher’s exact test. Font indicates significance and the fraction before each term shows the number of genes annotated with the GO term relative to the total number of such genes in the dataset. (b, top) A network plot of brain GO enrichment results was made in Cytoscape using EnrichmentMap (FDR Q value < 0.01, combined coefficients > 0.375, combined constant 0.5). Most highly connected nodes (b, bottom) are annotated using AutoAnnotate. Each node is a gene set and the size of each node represents the number of genes in the gene set. Edges (lines between nodes) represent overlap between gene sets and their width refer to the number of genes that are shared by the nodes. The color of each node represents enrichment scores (q-value).

### Single nuclei sequencing of the Orofacial Motor Cortex (OMC)

We assessed the quality of the data using cellranger (114 outliers removed, 3486 nuclei retained). For 3,486 nuclei that passed quality control, we did dimensional reductions (see Methods) and displayed them using t-distributed stochastic neighbor embedding (t-SNE; Figure 6a) (67) and uniform manifold approximation and projection (UMAP; Figure 6b) (68,69). Major brain cell types in 12 clusters were clearly identifiable based on canonical cell-type marker gene expression (Figure 6c). Major brain cell types in 12 clusters were clearly identifiable based on canonical cell-type marker gene expression (Figure 6c). We identified 5 clusters of nuclei as excitatory neurons, expressing high levels of Syt1 (synaptotagmin-1), two clusters as inhibitory neurons, which expressed high levels of Gad-2 (glutamate decarboxylase 2), one cluster of astrocytes, expressing Gfap (glial fibrillary protein), one cluster of oligodendrocytes, which expressed Mbp (myelin basic protein), and one cluster of endothelial nuclei, expressing high Vgfr (vascular endothelial growth factor receptor)(70–72). Normalized expression of the top ten marker genes for each brain cell type clearly distinguished the 12 nuclei clusters using t-SNE and UMAP (Figure 6d).

**Figure 6.**
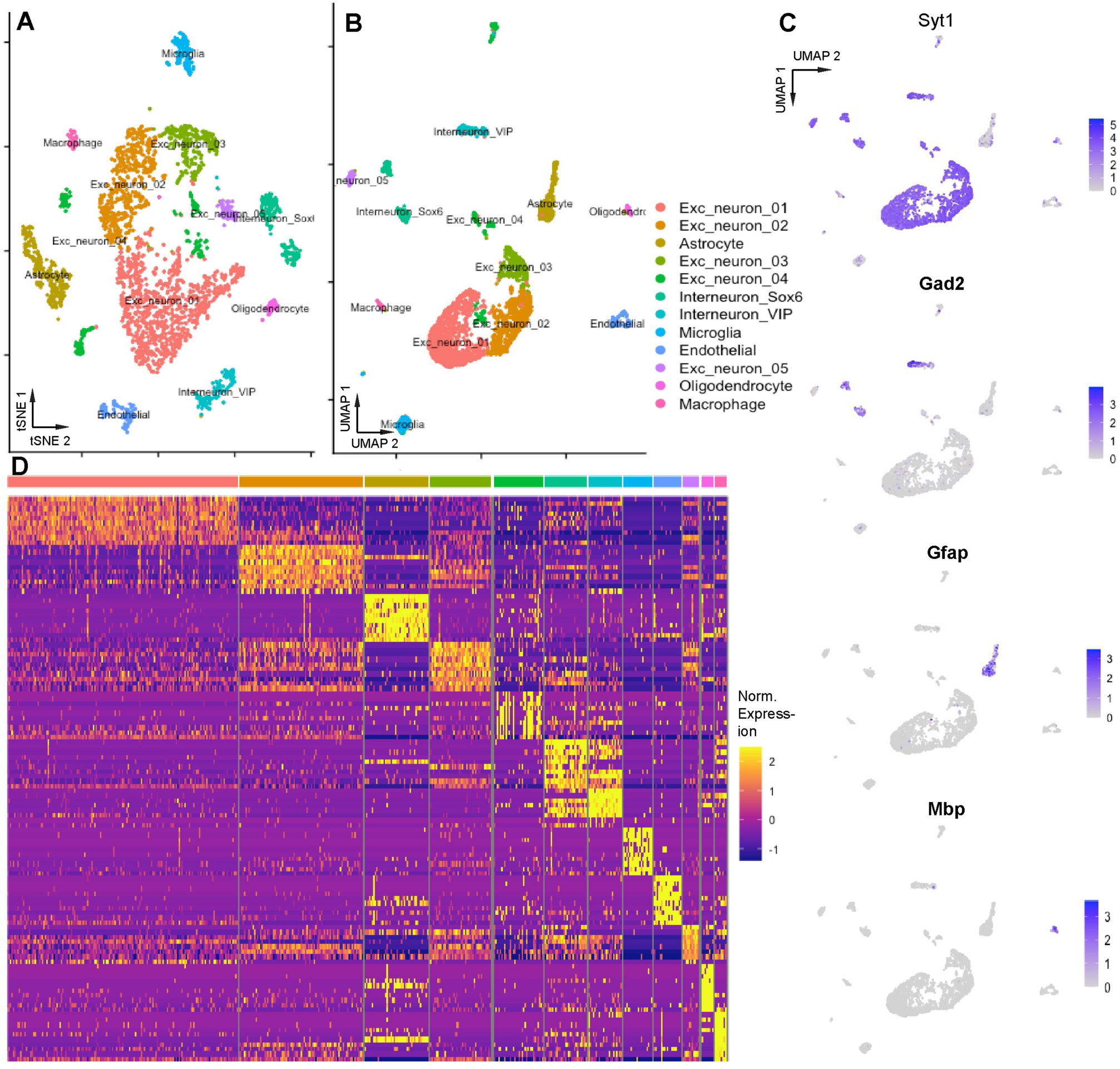
The *S. teguina* genome and transcriptome enable identification of brain cell types in a single-nuclei RNAseq dataset from brain area OMC. A. tSNE dimensional reduction of 3486 nuclei isolated from brain area OMC. B. UMAP dimensional reduction of same data C. Feature plots of normalized gene expression data for key brain cell type markers (Syt1: synaptotagmin-1, neuron; Gad2: glutamate decarboxylase 2, inhibitory neuron; Gfap: glial fibrillary protein, astrocyte; Mbp: myelin binding protein, oligodendrocyte; Vgfr: vascular endothelial growth factor receptor, endothelial cell). D. Heatmap of normalized gene expression data for the top 10 marker genes for each brain cell type identified.

## Discussion

We sequenced, assembled, and annotated a genome and transcriptome for Alston’s singing mouse, a model for complex social behavior and vocal communication. Transcriptome and gene annotation quality were validated using gene expression and functional enrichment analyses. Finally, we did single-nuclei sequencing of cells of the orofacial motor cortex (OMC), a region involved in vocal turn-taking in singing mice (31) and identified major cell types. The annotated genome and transcriptome will be a valuable resource that will allow characterization of the genetic basis of complex traits in singing mice as well as be useful for comparative studies more broadly.

### Genome and transcriptome assembly and annotation

By using three sequencing technologies, we were able to create a high quality *de novo* genome assembly. Short reads, like those generated by Illumina, provided the highest base-pair-level accuracy (73–75). Longer reads generated by PacBio’s SMRT sequencing (76,77), produced excellent contigs. Finally, contig scaffolding was facilitated by PromethION’s nanopore sequencing (78,79), which can sequence the longest stretches of DNA (80).

These sequencing efforts culminated in a 2.4 billion base pair genome. The size of the singing mouse draft genome is like that of other sequenced rodents such as house mice, *Mus musculus*, (strain: C57BL/6J, genome size: 2.5 billion bp) (81) and white-footed mice, *Peromyscus leucopus* (genome size: 2.45 billion bp) (82–84).

We assembled 754,907 transcripts into a *de novo* transcriptome with a contig N50 of 826. When aligned to the reference genome, 83.41% of the transcriptome mapped, indicating a quality transcriptome assembly (37,38). As expected, the contig N50 based on only the longest isoform per gene was lower than that of all transcripts since including all transcript isoforms can exaggerate N50 values.

We did differential expression and functional enrichment analyses to test the quality of the transcriptome and annotations. In support of our expectations for a quality assembly and annotation, we found tissue-specific gene expression profiles. Two distinct clustering methods showed that samples of the same tissues type, both technical and biological replicates, had identical (technical replicates) or very similar (biological replicates) expression patterns that differed greatly from other tissues. Functional enrichment analysis identified the putative function of differentially expressed genes across tissue type. Within the brain, we found enrichment of expected pathways such as synapse-related GO terms. A network-based approach supported these results, clustering related nodes into larger functional groups with brain-relevant annotations such as “glutamate neurotransmitter receptor.” This approach is useful because GO functional categories often share many genes and the results of GO analyses can often be redundant (53,55). A network approach clusters by gene overlap which allows the annotation of groups of similar gene sets, rather than annotating each gene set independently. The results of these analyses suggest that our transcriptome assembly is of high quality and the annotations we created are accurate. For our functional enrichment analyses, we focused on brain gene expression because we are interested in understanding how the nervous system drives complex behavior. A quality transcriptome assembly and gene annotations allowed us to examine gene expression of single nuclei to identify specific brain cell types that may contribute to behavior (see Single-nuclei sequencing of the OMC). Single-nuclei sequencing of regions implicated in song production (31,85) can be paired with other approaches such as sequence-based interventions and epigenetic profiling to understand how the brain patterns vocal output.

The genome, transcriptome, and annotation GTF file can be accessed at the UCSC genome browser (https://bit.ly/3hfUiIy). We identified 14,989 genes that have at 80% sequence similarity to reference genes. This is on the order of annotation efforts in other species (81,82,86,87). Although we used a sequence similarity approach, phylogenetic approaches may improve our ability to identify genes accurately, as sequence similarity does not necessitate shared function.

### Single-nuclei sequencing of the OMC

Dimensional reductions of the expression profiles of 3,486 OMC nuclei revealed 12 clusters that were categorized into major cell types based on marker genes. Most nuclei fell into 7 clusters that were categorized as neuronal, which we expected, since we used NeuN antibodies to enrich for neuronal nuclei prior to sequencing. Five of these clusters were identified as containing nuclei of excitatory neurons, expressing high levels of Syt1 (synaptotagmin-1), gene encoding a Ca2+ sensory for neurotransmitter release (72,88–90). Two inhibitory interneuron clusters were identified by expression of Gad-2 (glutamate decarboxylase 2), which encodes an enzyme that catalyzes the synthesis of GABA, an inhibitory neurotransmitter (72,91). Despite selecting for neuronal nuclei, a few small clusters of other cell types were also identified such as astrocytes (high Gfap) and oligodendrocytes (Mbp) (72). Clear identification of brain-cell types demonstrates the robustness of the singing mouse transcriptome and genome and demonstrates the broad applicability of 10X single cell/single nuclei technology to a nontraditional rodent species. The brain region we chose for single nuclei analysis is an important temporal regulator of the singing mouse advertisement song (31). By combining single-cell analysis of relevant brain regions with circuit-level studies (85), we can develop hypotheses about the role of each network node and the cellular mechanisms that underlie these functions. Together, these resources allow us to examine how the nervous system directs complex behavior.

### Uses of these resources

The contribution of a high-quality singing mouse genome and transcriptome increases the diversity of available model species and improves our ability to ask mechanistic questions. Singing mice are a particularly useful model for understanding how the brain drives complex behavior due to their unique, quantifiable phenotype, their tractability in the lab, and our ability to adapt existing neurobiological tools and resources for singing mice. The addition of genomic resources provides further opportunity to use singing mice to study novel questions in social cognition. For example, to explore the genomic basis of complex traits, we could use the genomic resources we have generated to examine gene regulation (e.g., ChIP-seq: (1); ATAC-seq: (93)), sequence evolution (e.g., tests of selection: (94–96)), and more. These data also contribute to a library of resources that can be used for larger comparative analyses. Increasing the diversity of model systems, through the addition of species that are well-suited to particular questions, is essential to understanding the mechanisms that drive complex traits.

## Abbreviations

PUREaCSF: Artificial cerebrospinal fluid
BLAST: Basic local alignment search tool
cDNA: Complementary DNA
DAPI: 4’,6’-diamidino-2-phenylindole
ENCODE: Encyclopedia of DNA elements
FITC: Fluorescein isothiocyanate
GDNA: Genomic DNA
GEM: Gel bead-in emulsion
GO MWU: Gene ontology Mann-Whitney U test
GTF: Gene transfer format
KEGG: Kyoto encyclopedia of genes and genomes
OMC: Orofacial motor cortex
PASA: Program to assemble spliced alignments
PCA: Principal components analysis
scRNASeq: Single-cell RNA sequencing
STAR: Spliced transcripts alignment to a reference
tSNE: t-distributed stochastic neighbor embedding
UMAP: Uniform manifold approximation and projection for dimension reduction

## Declarations

### Ethics approval and consent to participate

All experimental procedures were approved by the University of Texas Austin and New York University Grossman School of Medicine IACUC. All methods were done in accordance with IACUC regulations. We confirm that we have reported this research following the ARRIVE guidelines for the reporting of animal research.

### Consent for publication

Not applicable.

### Availability of data and materials

Sequence and annotation data are available at the UCSC Genome Browser (https://bit.ly/3hfUiIy). Single-cell sequencing data is accessible by request.

### Authors’ Contributions

SMP, AH, SB, and MAL designed the research. AH and MO generated data. PWF generated scRNAseq data and analyzed the single nuclei expression data. CM, AK-J, SB, PZ, and SKS analyzed genome and bulk transcriptome data. SKS wrote the manuscript. SKS, PWF, and AK-J prepared figures and tables. All authors read and approved the final version of the manuscript.

## Acknowledgements

We would like to thank the Genome Technology Center (GTC) and UT Genomic Sequencing and Analysis Facility (GSAF) for expert library preparation and sequencing. We are grateful to the Applied Bioinformatics Laboratories (ABS) for providing bioinformatics support and helping with the analysis and interpretation of the data. GTC and ABL are shared resources partially supported by the Cancer Center Support Grant P30CA016087 at the Laura and Isaac Perlmutter Cancer Center. This work has used computing resources at the NYU School of Medicine High Performance Computing (HPC) Facility and the Texas Advanced Computing Center (TACC).

